# Generalizing Genetic Risk Scores from Europeans to Hispanics/Latinos

**DOI:** 10.1101/242404

**Authors:** Kelsey E. Grinde, Qibin Qi, Timothy A. Thornton, Simin Liu, Aladdin H. Shadyab, Kei Hang K. Chan, Alexander P. Reiner, Tamar Sofer

**Affiliations:** Department of Biostatistics, University of Washington, Seattle, WA, USA; Department of Epidemiology & Population Health, Albert Einstein College of Medicine, Bronx, NY, USA; Department of Epidemiology, Brown University, Providence, RI, USA; Department of Family Medicine and Public Health, University of California San Diego, San Diego, CA, USA; Division of Public Health Sciences, Fred Hutchinson Cancer Research Center, Seattle, WA, USA; Division of Sleep and Circadian Disorders, Brigham and Women’s Hospital, Boston, MA, USA; Department of Medicine, Harvard Medical School, Boston, MA, USA

**Keywords:** genetic diversity, admixed populations, linkage disequilibrium

## Abstract

Genetic risk scores (GRSs) are weighted sums of risk allele counts of single nucleotide polymorphisms (SNPs) associated with a disease or trait. Construction of GRSs is typically based on published results from Genome-Wide Association Studies (GWASs), the majority of which have been performed in large populations of European ancestry (EA) individuals. While many genotype-trait associations have been shown to generalize from EA populations to other populations, such as Hispanics/Latinos, the optimal choice of SNPs and weights for GRSs may differ between populations due to different linkage disequilibrium (LD) and allele frequency patterns. This is further complicated by the fact that different Hispanic/Latino populations may have different admixture patterns, so that LD and allele frequency patterns may not be the same among non-EA populations. Here, we compare various approaches for GRS construction, using GWAS results from both large EA studies and a smaller study in Hispanics/Latinos, the Hispanic Community Health Study/Study of Latinos (HCHS/SOL, *n* = 12, 803). We consider multiple ways to select SNPs from association regions and to calculate the SNP weights. We study the performance of the resulting GRSs in an independent study of Hispanics/Latinos from the Woman Health Initiative (WHI, *n* = 3, 582). We support our investigation with simulation studies of potential genetic architectures in a single locus. We observed that selecting variants based on EA GWASs generally performs well, as long as SNP weights are calculated using Hispanics/Latinos GWASs, or using the meta-analysis of EA and Hispanics/Latinos GWASs. The optimal approach depends on the genetic architecture of the trait.

## Introduction

Genetic Risk Scores (GRSs) summarize the genetic component of a disease or quantitative (continuous) trait and are typically constructed as the weighted sum of risk alleles of single nucleotide polymorphisms (SNPs) associated with the trait of interest. GRSs are routinely used in public health genetics research for a wide range of applications, such as improving disease and trait prediction;^1^ studying the shared genetic basis between traits;^2^ increasing power by integrating over multiple variants rather than one variant at a time; and Mendelian Randomization studies,^3^, ^4^ in which a GRS associated with one trait is used as an instrumental variable in a causal analysis of the association of the trait with another outcome. SNPs and weights used to construct GRSs are usually selected based on findings from published studies, such as Genome-Wide Association Studies (GWASs).^5^ However, most GWASs to date have been performed in studies of individuals of exclusively or predominantly European genetic ancestry (EA). This poses a difficulty for GRS construction in non-EA populations.

Using EA GWAS to select SNPs and choose weights for GRSs in non-EA populations seems like a reasonable approach, in particular, Hispanics/Latinos are admixed with European ancestry, and previous studies have shown that many EA GWAS results generalize to Hispanics/Latinos.^7, 9^. Furthernore, the EA GWASs are very large (with tens or hundreds of thousands of individuals), therefore having large statistical power to detect the most strongly associated variants from genomic association regions and obtain precise estimates of effects sizes. In fact, Dudbridge (2013)^6^ studied power and prediction accuracy of GRSs and suggested that, for prediction (rather than association testing), hundreds of thousands of subjects may be needed to estimate SNP effects. While such numbers are available in EA GWASs, they are not currently available in GWASs of diverse populations such as Hispanics/Latinos. For example, in the Hispanic Community Health Study/Study of Latinos (HCHS/SOL), a large cohort study of Hispanics/Latinos, there are fewer than 13,000 individuals who consented for genetic studies. Of these, about 7,000 individuals participated in a GWAS of diabetes,^7^ which is the largest published GWAS to date in Hispanics/Latinos. In contrast, the largest published GWAS of diabetes^8^ meta-analyzed multiple studies of European ancestry (*N* = ~ 70,000 cases and controls), and a smaller number of population studies of other ancestries (including about 2,500 Mexicans).

There are a number of drawbacks to using EA GWAS for GRS construction in non-EA populations. Specifically, linkage disequilibrium (LD) patterns vary across populations,^10^ rendering different best available tag SNPs between populations; allele frequencies often differ across populations; and, at least for some traits, effect sizes differ between populations and allelic heterogeneity exists.^11^ Admixed populations, such as Hispanics/Latinos, may have different genetic architecture and effect sizes at a genetic association region compared to an ancestral population due to gene-gene (epistasis) or gene-environment interactions, or because a causal variant monomorphic in one ancestral population.^11^, ^7^ For example, Belsky et al. (2013)^12^ constructed a GRS for obesity based on EA GWAS results, and found that its utility for an African American (AA) population was low, and much lower than that for an EA population. Martin et. al. (2017)^13^ studied transferability of GRSs constructed based on single-ancestry GWASs to other ancestries, and demonstrated that scores inferred from EA GWASs may perform poorly in other ancestries. Collectively, these studies highlight the need to adapt GRSs construction methods to diverse ancestries.

How to best construct a GRS for a study in a Hispanic/Latino population is still an open question. Should one use only the information published in a large, primarily EA study? or can we use results from a smaller, non-EA study? In particular, will incorporating information from these lower-powered (smaller sample size) non-EA GWAS in fact improve GRS construction, or will it instead introduce harmful variability? Here, we take a systematic, empirical approach to constructing GRSs for Hispanics/Latinos, based on GWAS results from large population studies of European ancestry and medium-sized studies of Hispanics/Latinos. We use published EA GWASs results and GWAS findings from the HCHS/SOL to construct and evaluate GRSs in an independent study of Hispanic/Latina women from the Women Health Initiative (WHI). We support our results with simulations mimicking potential genetic architecture within a single, trait-associated genomic region.

## Materials and Methods

Let *y_i_* be a quantitative trait measured on the *i*th participant, *i* = 1,…,*n*, and *x_i_* a *k* × 1 vector of covariates such as confounders. Let *g*_1_*_i_*,…,*g_Pi_* be allelic counts or dosages of *P* independent variants associated with *y_i_*, and *α*_1_,…,*α_P_* their effect sizes, so that the additive linear model holds:

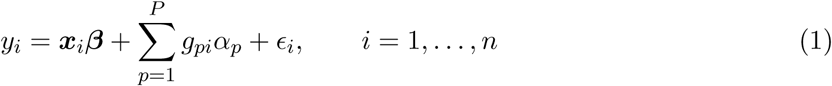
where *ϵ_i_* are residual errors. An optimal GRS for *y_i_* is 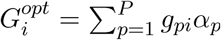, the weighted sum with the causal genotypes and their true effects.

### Issues in selection of SNPs for GRS

In reality, we do not know which are the true causal genotypes for a trait, so we have to select a set of SNPs to use in our GRS. Often, the data we have at our disposal for selecting SNPs are derived from a genotyping platform that did not interrogate all sequence genotypes, but rather a reduced set of a few million (or fewer) variants. For GRS construction, we often have only a set of associated genotypes that likely tag a subset of the causal genotypes.

Let *g_p_* be a causal genotype in populations of European ancestry, and let 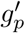 be a tag SNP to *g_p_* that was detected in an association study, perhaps because *g_p_* was not genotyped at all. It is well known^14^ that the size of trait association at 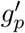 is related to the LD of 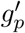 with *g_p_*, denoted by *ρ_p_*, so that 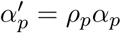. Given a large enough data set, we expect that the lead variant (the variant with strongest association in the region) will be the one with |*ρ_p_*| closest to the maximal value 1, among all available (genotyped or imputed) variants.

Further complicating this situation is the fact that tag SNPs may differ across populations. Assume the simple scenario of a single causal SNP in an association region, two ancestral populations *P*_1_ and *P*_2_, the same effect size in all populations. Also, assume that in the admixed population (ADM) the proportion of genomic intervals containing the causal variant inherited from populations *P*_1_ and *P*_2_ is *a* and (1 − *a*), respectively. This is demonstrated in Figure 1, which shows that even if the same tag SNPs were available in the two ancestral populations *P*_1_ and *P*_2_, and we knew which tag SNPs were the best in each population, it is not clear which is the best tag SNP in ADM. This becomes even more complicated when there are multiple ancestral populations, when SNP availability differs due to different genotyping platforms, and when effect sizes differ between ancestral populations.

**Figure 1:**
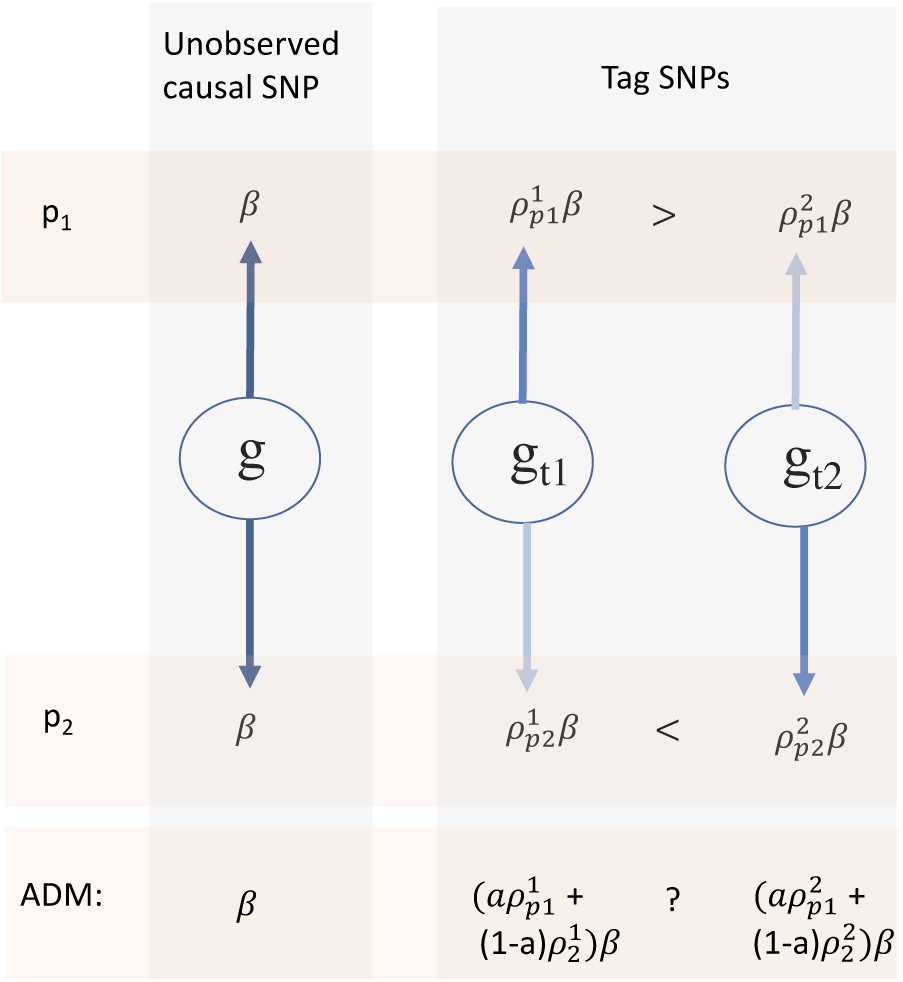
Tag SNPs likely have different linkage disequilibrium patterns with the causal SNP *g* in populations *P*_1_ and *P*_2_, so that different observed tag SNPs may have stronger associations with the trait in the two populations. In the admixed population (ADM), the associations at the the tag SNPs depend on *a*, the proportions of chromosomes with this genomic regions inherited from the two ancestral populations.

In simulations studies, we investigated the impact of genetic architecture by focusing on a single genomic region at a time, while in data analysis we considered the overall results of systematic genome-wide approaches for GRS construction.

### GRS construction

#### SNP selection

Consider a GWAS performed in a study of European ancestry. Association results are available for *d* variants, with *p*-values 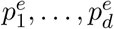 and effect size estimates 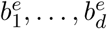 Assume that almost all variants are also available in a GWAS of individuals with admixed ancestry, such as the HCHS/SOL, with corresponding *p*-values and effect sizes 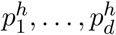, and 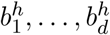. Based on the information from both EA and HCHS/SOL GWASs, we can perform fixed-effects meta-analysis (META) to produce meta-analytic *p*-values and effect sizes 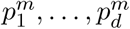, and 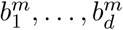. We can also perform generalization analysis^10^, to test the composite null hypothesis that is rejected if an association exists in both the EA population and in the HCHS/SOL study, and get *r*-values for these variants 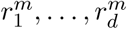.

We created lists of candidate SNPs for GRSs by filtering variants based on these measures (**p***^e^*, **p***^h^*, **p***^m^*, **r***^m^*) and with varying thresholds. We considered the *p*-value thresholds 5 × 10^−8^, 1 × 10^−7^, 1 × 10^−6^, 1 × 10^−5^, 1 × 10^−4^, 1 × 10^−3^ and *r*-value thresholds 0.05, 0.1, 0.2, 0.3, 0.4, 0.5. For generalization analysis, we initially took all variants with *p*-value < 10^−6^ in the EA GWAS, then performed generalization analysis using these SNPs to compute *r*-values. Therefore, by construction, smaller lists of SNPs are considered using this approach.

After subsetting the list of available SNPs by the selection criteria (e.g., all SNPs with *p*-value smaller than 10^−6^ computed in the EA GWAS, all SNPs with *p*-value smaller than 5 × 10^−8^ in the meta-analysis GWAS, etc.) to generate a list of candidate SNPs, we further pruned the list to keep only the lead SNP from every genomic region of 1Mbp. The first lead SNP is defined as the most significant SNP in the list; then, all SNPs in the 1Mbp region centered at this SNP are removed from the list of candidate SNPs. We continued this process of identifying the next lead SNP and removing SNPs in its proximity until no SNP remained. We also considered another method for selecting SNPs, which we call “matched SOL”. For this approach, we took the list of lead SNPs from the EA GWAS, and searched for the lead HCHS/SOL SNP among the common (MAF ≥ 0.05) genotyped SNPs within 1Mbp around it.

In simulation studies, where we focus on a single association region, we did not filter variants. Instead, we selected only a single lead SNP from the region based on the same criteria described above (lead EA, lead in meta-analysis, etc.).

#### SNP weights

The optimal GRS weights reflect the true size of association between each SNP in the GRS and the trait. In practice, the size of this association must be estimated. We considered the effect size estimates computed in the EA GWAS, which may be very accurate in the EA population but potentially less appropriate in the admixed population due to different LD patterns in the two populations. Therefore, we also considered the effect size estimated in an admixed population (HCHS/SOL in the data analysis) and the effect size estimated in fixed effects meta-analysis. We also compared these to no weights (or *α_p_* = 1 ∀*p*) because unweighted GRS are often used in practice.^7^, ^2^

#### Evaluating the GRS

To evaluate GRS approaches, we constructed GRSs in an independent validation data set based on SNP selection and weights computed in the training data set. In simulations, we used a validation data set with different admixture patterns than the training data set, to reflect the fact that different admixed populations often differ to some extent in their admixture patterns. In our data analysis, the training data set was HCHS/SOL and the validation data set was the WHI SNP Health Association Resource (SHARe) Hispanic/Latina women.

Let the GRS for participant *i* in the validation data set be

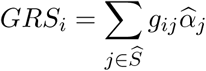
where 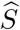 is the selected set of SNPs (which is likely different than the true causal set *S*, and in the simulation has a single SNP only), and 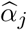 is the estimated effect of the *j*th SNP is the set. We considered two measures for evaluation. For simulation studies we used the Mean Squared Prediction Error (MSPE), computed by

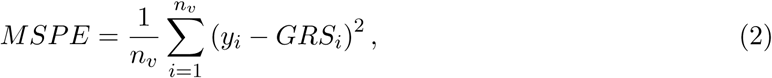
across the *n_v_* individuals in the validation data set, where we report the squared root of the MSPE (RMSPE). In data analysis, we computed the variance explained by each GRS in a regression model adjusted for sex, age, and the first five principal components (PCs) of genetic ancestry. This was calculated by first fitting a model with these covariates, but without the GRS, and obtaining the residual variance denoted by 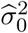, then fitting a model that also included the GRS and obtaining the residual variance 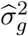. The estimated percent variance explained is 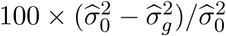.

### Simulation study

Our simulation studies focused on the impact of LD and variability in the estimation of effect sizes (due to sample size and admixture) on GRSs. We simulated genotypes in a 1Mbp genomic region for a large European sample (*n_EA_* = 50, 000), a moderately sized admixed sample (*n_ADM_*_12_ = 12, 000), and a small admixed sample (*n_ADM_*_5_ = 5, 000). ADM_12_ and ADM_5_ are admixed populations (details below) with the same two ancestral populations—CEU (European ancestry) and YRI (African ancestry)—but different admixture proportions—20% or 40%. We simulated quantitative traits under a few potential genetic architectures, assuming one or two causal SNPs, that are either shared or different between the two ancestral populations, in the region. Each simulation setting was repeated 500 times. Details about simulating admixed populations genomic association regions and architectures, are provided in the Supplemental Information.

### Evaluating similarity between LD patters

We evaluated the similarity between LD patterns of the simulated populations CEU, ADM_0.2_, and ADM_0.4_. For each SNP *j* = 1,…, 617 in the simulated genomic region, we identified its best tag SNP in CEU, ADM_0.2_, and ADM_0.4_ by finding the SNP *j*′ ≠ *j* which had the highest LD (*r*^2^) with SNP *j* in that sample. Given a population of interest ADM_0.2_ (ADM_0.4_), we calculated the proportion of SNPs in the region had a tag SNP with higher LD in CEU compared to ADM_0.4_ (ADM_0.2_), and the other way around.

### Using the HCHS/SOL to develop Hispanic/Latino specific GRSs

#### Previously published EA GWASs and traits

We considered three groups of traits with previously published GWAS results: anthropometric, comprising of height, body mass index (BMI), hip circumference (HIP), waste circumference (WC), and waste-to-hip ratio (WHR) from the GIANT consortium GWAS;^18–20^ blood pressure traits, comprising of systolic and diastolic blood pressure (SBP, DBP), pulse pressure (PP = SBP-DBP), and mean arterial pressure (MAP = DBP+1/3PP), with GWASs performed by the International Consortium of Blood Pressure (ICBP)^21^, ^22^; and finally, blood count traits, including white blood cell count (WBC), platelet count (PLT), and hemoglobin count (HGB).^23–25^ Note that for these blood count GWASs, we used just the EA GWAS results and not the trans-ancestry results that were also available.

#### The Hispanic Community Health Study/Study of Latinos (HCHS/SOL)

The HCHS/SOL is a community-based cohort study following 16,415 self-identified Hispanic/Latino participants with initial visits between 2008 and 2011^26^. Participants were recruited into the study in four field centers (Chicago, IL, San Diego, CA, Bronx, NY, and Miami, FL) via a two-stage sampling scheme, by which community block units were first sampled, followed by households within the block units. Some or all household members were recruited. The sampling probabilities were set preferentially towards sampling Hispanics/Latinos.^27^ In total, 12,803 study participants consented to genetic studies. Henceforth, we focus on this subset when describing the HCHS/SOL population. The HCHS/SOL participants are very diverse, and usually self identify as belonging to one of six background groups: Central American, Cuban, Dominican, Mexican, Puerto Rican and South American. Genotyping, imputation, and quality control in the HCHS/SOL have been described elsewhere.^28^

#### Genome-Wide Association Studies in the HCHS/SOL

All HCHS/SOL analyses adjusted for sex, age, log-transformed sampling weights (to prevent potential selection bias due to the study design), and the first five principal components reflecting ancestry (to control for population stratification). Analyses used mixed models with variance components due to genetic relatedness, household, and block unit sharing. GWASs for blood count traits in the HCHS/SOL^29^, ^30^, ^11^ used imputation based on the 1000 Genomes Project^31^ phase 1 reference panel. For blood pressure (BP) traits, stratified analyses were performed,^32^, ^33^ with stratification by Hispanic/Latino background groups (Central American, Cuban, etc.). We used results from the meta-analysis across all background groups as **p***^h^*, **b***^h^* for GRS construction. The GWASs for these traits used the phase 3 1000 Genomes Project reference panel. Anthropometric traits GWASs in the HCHS/SOL all used phase 1 imputation of the 1000 Genomes Project, but the height GWAS was stratified by background group, while the BMI, WHR, WC and HIP GWAS were not. The height GWAS was further adjusted for the status of being born in the US.

#### The Women’s Health Initiative SNP Health Association Resource (WHI SHARe)

The WHI is a long-term health study of postmenopausal women in the U.S. A total of 161,808 postmenopausal women aged 50-79 years old who were recruited from 1993 through 1998 and from 40 clinical centers throughout the United States.^34^ Ten of the 40 WHI clinical centers with expertise and access to specific minority groups (American Indian, black, Asian American or Pacific Islander, and Hispanic) were selected to serve as minority recruitment sites. Clinical information was collected by self-report and physical examination. A total of 5,469 self-identified Hispanic Americans (HA) were consented to genetic research and were eligible for WHI-SHARe. Due to budget constraints, we randomly selected a subsample of 3,642 (66.6%) HA women. DNA was extracted by the Specimen Processing Laboratory at the Fred Hutchinson Cancer Research Center from specimens that were collected at the time of enrollment. All participants provided written informed consent as approved by local human subjects committees. Genotype data are from the Affymetrix Genome-Wide Human SNP Array 6.0 that contains 906,000 single nucleotide polymorphisms (SNPs) and more than 946,000 probes for the detection of copy number variants. The genotype data were processed for quality control, including call rate, concordance rates for blinded and unblinded duplicates, and sex discrepancy, leaving 871,309 unflagged SNPs with a genotyping rate of 99.8% and 3586 HA women used in the current analysis. Genotype Imputation was carried out with MaCH. For imputation in HA samples, we used reference panel of HapMap III CEU + MEX (Mexican ancestry in Los Angeles, California) + YRI samples for a total of 1,387,466 SNPs (MAF > 1%), of which 1,368,178 SNPs met the quality threshold of *r*^2^ > 0.3.^35^

## Results

### Simulations

Our simulations focused on a single, 1Mbp genomic region, which contains 617 SNPs. We first evaluated the similarity of LD patterns between potential reference populations (CEU, ADM training data set) and the test data set, and then evaluated the combined impact of LD, genetic architecture, and variability in effect estimates on GRS performance.

### LD patterns in the admixed samples are more similar to each other than to CEU

Table 1 reports the proportion of SNPs in our locus for which the best tag SNP in CEU had higher LD with the causal SNP in the admixed validation data set, compared to the LD of the best tag SNP in the training admixed test data set. The tag SNPs in CEU and the training ADM population were often the same (46-50% of the time). When they differed, the tag SNP identified in the training admixed population was usually a better tag of the causal SNP in the test admixed population.

**Table 1:** The percentage of SNPs in the investigated locus for which the tag SNP in the population referenced in the table column had higher LD with the causal SNP in the population referenced in the table row. When the test population is ADM_0.2_ (ADM_0.4_), the training ADM population is ADM_0.4_ (ADM_0.2_).

| Test population | CEU better tag | Training ADM better tag | Same tag |
| --- | --- | --- | --- |
| ADM<sub>0.2</sub> | 4% | 46% | 50% |
| ADM<sub>0.4</sub> | 9% | 45% | 46% |

### Performance of GRSs in the simulated test data sets

Our simulation study considered four different scenarios of genetic architecture at the locus, focusing only on the effect of LD and sample sizes, and assuming that effect sizes are the same for all causal SNPs in populations. Each simulation included multiple, distinct choices of causal SNP(s). Details are provided in the Supplemental Information. For each choice of causal SNP(s) we repeated the simulation 500 times, constructing 12 GRSs each time (all combinations of the four selection and three weight estimation approaches), and we recorded the median MSPE for each GRS across the 500 repetitions.

Results are visualized in figures depicting the distributions of the medians squared-root MSPEs (RMSPEs) across the various choises of causal SNPs in each of the simulation scenarios. Figure 2 provides these distributions for two simulation scenarios, when ADM_12,0.2_ and CEU were used as a training data sets and ADM_5,0.4_ was the test data set. Figures corresponding to all other scenarios and settings are provided in the Supplemental Information.

**Figure 2:**
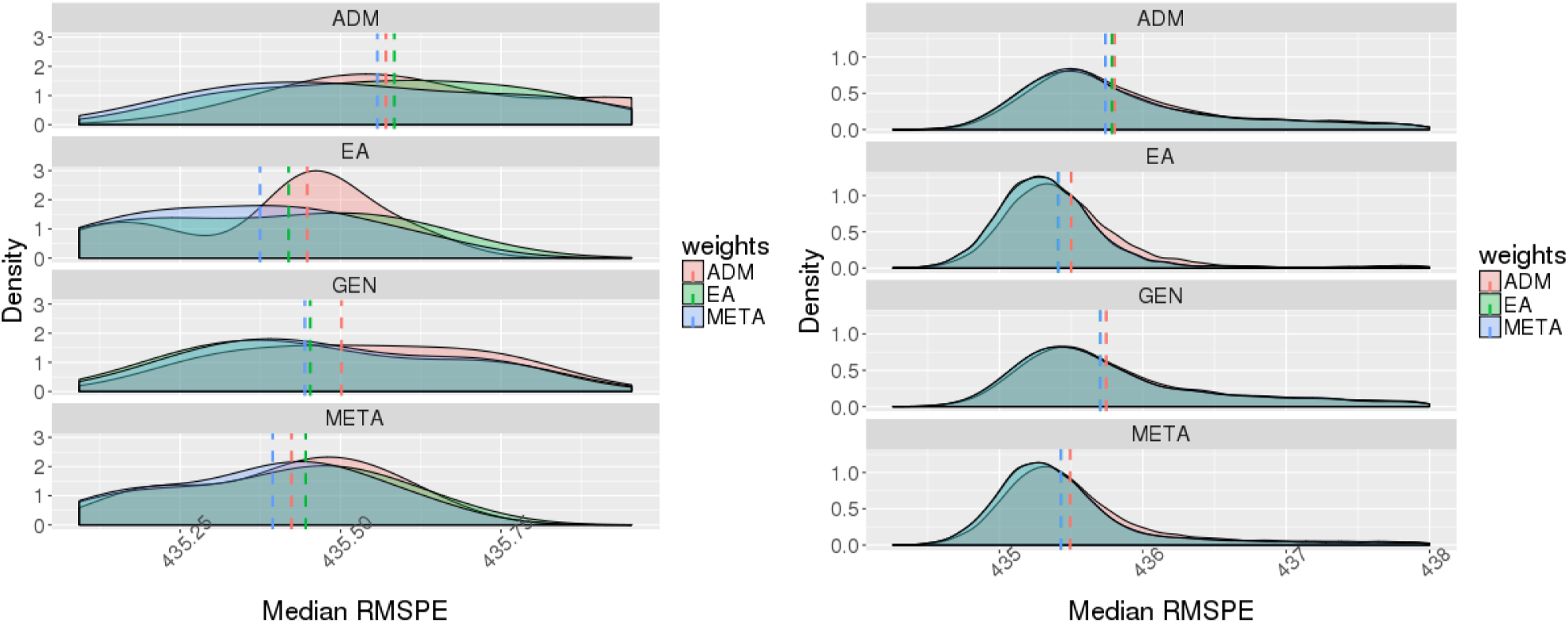
The smoothed distribution of median RMSPEs, where each median was computed over 500 repetitions of the simulations for a single choice of causal SNP(s), and the distribution is across all possible choices of causal SNP(s) in the same simulation scenario. The left panel corresponds to scenario 2 (a single causal SNP, which is monomorphic in YRI), and the right panel corresponds to scenario 3 (two causal SNPs, one of them is monomorphic in CEU). The training data sets were EA and ADM_12,0.2_, while the test data set was ADM_5,0.4_. Dashed vertical lines correspond to median of the plotted distribution. In the right panel, the lines corresponding to EA and META weights often overlap.

In almost all simulation scenarios, SNP selection by ADM training data set and weights calculated by ADM (for all other SNP selections) performed the worst. Only in simulation scenario 2, in which there are two causal SNPs with one being polyorphic only in YRI, computation of weights in the ADM training data is sometimes advantageous over EA weights. However, this was true only when ADM_12,0.2_ was the training data set, but not when ADM_12,0.4_ was the training data set. Other than that, both EA and META SNP selections and weights constructions usually performed similarly, with a few more settings in which META weights outperformed EA weights. We do note that all types of GRS suffered from outlying scenarios: specific combinations of causal SNP(s) in which one GRS produced extremely large RMSPEs. However, on average we see better performance of GRSs based on the META and EA GWAS relative to the other two selection approaches when the causal SNPs’ effect sizes are the same across populations.

### Performance of GRSs in the Women’s Health Initiative data set

We constructed GRSs for 3,642 Hispanic/Latina women from the WHI based on EA GWAS, HCHS/SOL GWAS, and combinations of the two for each of 12 traits. The results differed considerably across families of traits (anthropometric, blood count, and blood pressure), so we focus on each separately. Figures comparing the percent of variance explained by each GRS for each trait are provided in the Supplementary Information. Here, Figure 3 illustrates the patterns observed in each trait family by comparing the variance explained by GRSs for one trait from each family (BMI, HGB, and MAP). Figure 3 compares the performance of GRSs that selected SNPs based on EA GWAS, but with various choices of *p*-value thresholds and weights.

**Figure 3:**
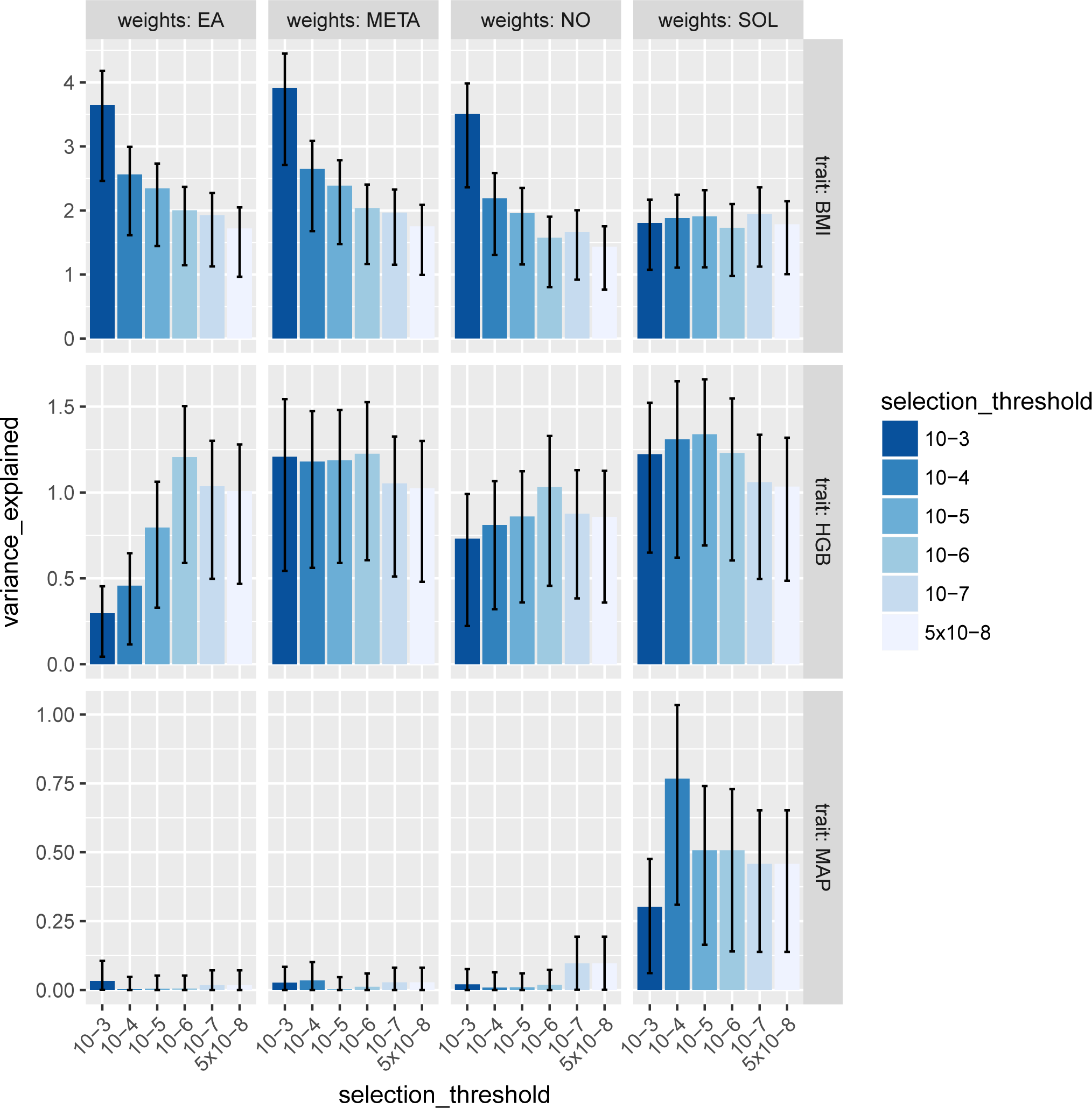
The percent of variance explained by GRSs for body mass index (BMI), hemoglobin count (HGB) and mean arterial pressure (MAP) in the WHI Hispanic/Latina women. For all GRSs in this figure, SNPs were selected based on their *p*-values in the EA GWAS, with various thresholds considered (marked by color), and pruned so that no two selected SNPs were closer than 500Kbp. SNP weights were computed by one of four methods (corresponding to columns): *EA* = estimated effect sizes in the EA GWAS, *META* = estimated effect sizes in the meta-analysis of the EA and HCHS/SOL GWASs, *NO* = unweighted GRS, and *SOL* = estimated effect sizes in the HCHS/SOL GWAS. Error bars indicate 95% confidence intervals for the percent of variance explained, based on 1,000 bootstrap samples from the WHI data set.

In general, our data analysis demonstrates that selecting variants for the GRS based on association results in the EA GWAS performs well, in agreement with the simulation results. The best choice of weights is usually estimated effect sizes from the HCHS/SOL GWAS or from the metaanalysis of the EA and HCHS/SOL GWASs (META). We now summarize the results by families of traits.

### Anthropometric traits

We considered five anthropometric traits: height, BMI, WC, WHR and HIP. Across all traits in this family, selecting SNPs using the EA GWAS or META performed similarly well, except in the case of WHR where selection based on EA has the slight advantage. The best choice of weights was also the same across all anthropometric traits, with weights from the EA GWAS and the meta-analysis GWAS performing similarly well. Optimal *p*-value thresholds for selecting SNPs differed across traits: for BMI, HIP, WC, and WHR the highest performing GRSs usually started from the list of SNPs with *p*-value < 10^−3^, while for height the optimal threshold was 10^−5^ − 10^−6^. The best performing GRSs explained about 10% of the variance of height in the WHI, 3% of the variance of BMI and HIP, 2.5% for WC, and less than 2% for WHR.

### Blood count traits

For blood count traits (PLT, WBC, HGB), the best performing GRSs were constructed either by selecting SNPs with EA GWAS, or META, *p*-values < 10^−5^ − 10^−3^ (optimal threshold varied by trait). For two traits (HGB and PLT), SNP selection using generalization also performed very well. The best performing weights were either the estimated effect sizes from HCHS/SOL (PLT, WBC) or META (HGB). Variance explained for the best GRSs ranged from ~1.3% (HGB) to 4% (PLT).

### Blood pressure traits

In contrast to the other trait families, for blood pressur traits (SBP, DBP, PP, MAP) selection of variants using the EA GWAS results performed poorly—unless weights were the effect size estimates from the HCHS/SOL GWAS, in which case the GRSs performed very well in comparison to other options. On the other hand, when selecting variants according to META, the GRS performance was robust to the choice of weights. Still, overall the GRSs that explained the highest percentage of variance used HCHS/SOL-based weights. An exception was the trait PP, for which selecting lead SNPs based on generalization analysis performed best, and was not sensitive at all to weight computation. However, the total percent of variance of PP explained by the best GRS was quite low: 0.4%. In comparison, the BP trait with the highest percent of variance explained by a GRS was MAP (1% explained by the best GRS). The optimal *p*-value threshold for SNP selection varied among traits and selection approaches.

## Discussion

We studied several approaches for constructing GRSs in Hispanic/Latino populations, using GWAS results from independent studies in large populations of European ancestry (EA) and medium-sized GWASs in Hispanics/Latinos. We studied the performance of GRSs constructed using these approaches on an independent data set. Results differed by trait. For anthropometric traits, the best strategies for both selecting variants for the GRS and for computing weights were based on either the EA GWAS or the meta-analysis of GWAS results from EA and HCHS/SOL, with a slight, but possibly negligible, advantage to selection based on meta-analysis results. For blood count traits (hemoglobin, white blood cell, and platelet counts), there was a slight advantage for selecting SNPs based on meta-analysis results, and it was generally preferable to use weights based on the HCHS/SOL GWAS. For blood pressure traits (SBP, DBP, MAP and PP), the best strategy overall was to select SNPs using the EA GWAS results and use weights from the HCHS/SOL GWAS, although the GRSs were more robust to different calculations of weights when meta-analysis results were used to select SNPs.

Our simulation studies were performed in simplified scenarios in which the effect sizes were the same in both ancestries, and in the admixed populations themselves. The LD patterns between two admixed populations with different admixture proportions were more similar to each other than the LD patterns between an admixed population and the simulated EA population, and further, LD estimates were very precise, with the same *r*^2^ observed when using 5,000 or more individuals for estimation. Therefore, given large enough sample size we expect that one admixed population will be a better reference for GRS construction in another admixed population compared to an EA population, and these simulations demonstrate how sample sizes used in the reference GWAS impact consequent GRS construction under various genetic architecture settings. In simulation results, usually the GRSs with SNP selection using EA GWAS and weights using META performed best, and weights using the admixed training data sets usually performed poorly. This results is supported by other work^6^ that suggested that many thousands of individuals are required to effectively calculate effect sizes to be used as SNP weights.

In the data analysis, however, in some cases GRSs weighted by the HCHS/SOL GWAS where superior to other weights. One possible explanation to the improved performance of the HCHS/SOL weights in data analysis is a strong signal-to-noise ratio for some traits and SNPs. Another possible reason is that the simplifying assumptions of the simulations do not hold and for some traits the SNP effect sizes differ by ancestry, or by population (e.g. due to gene-environment interaction). Thus, if the EA weights are inadequate despite the large sample size, it suggests that the estimates themselves are not appropriate for the admixed population. This is in agreement with the work of Coram et al.,^36^ who assumed different effect sizes between populations, and found that estimating effect for risk prediction purposes is useful in ethnically-matched population, while SNP selection using EA GWAS is generally appropriate.

Our study has a few limitations. First, we looked only at the performance of GRSs in independent validation data sets, so our results do not inform the construction of GRSs to be used in the same study (e.g., a Mendelian Randomization study). Furthermore, the independent validation study in our data analysis, WHI, only includes female participants, while our training studies, EA GWAS results and the HCHS/SOL, included both males and females. As gene-sex interactions likely exist, the GRSs constructed using the general population may not be optimal for women. However, this is unlikely to introduce any systematic biases to the SNP selection and SNP weights calculation procedures, so the relative performance of the GRS construction approaches should not be impacted. We did not investigate the entire literature for each trait and then investigate each of those loci separately, as is sometimes done in practice, but instead applied the same algorithm to each trait, based on two reference GWASs. While the first approach is useful for investigators who work with a single GRS and want to optimize it, it is also more case-dependent, and less generalizable. Our systematic approach is easier to apply on a number of traits and is appropriate for drawing general conclusions. Finally, we did not use multi-ethnic GWASs for GRS construction, but rather focused on EA and and Hispanic/Latino GWASs. Our goal was to more clearly delineate properties of the genetic architecture similarities or differences between populations. Another limitation of the current study is the lack of systematic investigation into generalizability of our results to other types of populations and varying sample sizes. It is a topic of future research as results from larger diverse studies become available. In the Supplementary Information, we report the results of a secondary analysis repeating the same data analysis reported in the manuscript, while evaluating GRSs on WHI African American women. Interestingly, the pattern of results is generally similar. This suggest that leveraging trans-ethnic information into GRS construction is beneficial.

Other recent methodological work on GRSs has been performed primarily in the context of EA populations. Shi et. al. (216)^37^ suggested to penalize the estimated effect sizes used in a GRS, by fitting an *ℓ*_1_-penalized regression. It is a topic of future work to suggest an approach that reduces the computational burden of applying shrinkage estimation procedure in mixed models and test its utility for improving the effect size estimates in an admixed population training data set. Vilhjálmsson et. al. (2015)^38^ proposed LDpred for incorporating information from GWAS summary statistics and a reference panel to use information from multiple SNPs, rather than only the lead SNP, from an association region. While they demonstrated this method to be useful under specific priors for genetic architecture, their approach hinges on having a good reference panel. Different admixed populations differ in their admixture patterns, so the same reference panel may not be appropriate across the board. It will be interesting to study and potentially extend this^38^ approach to admixed populations, despite the lack of training and testing samples with the same LD structure.

## Supplemental Data

The Supplemental Data includes detailed information about the simulation studies and figures with results from simulations, and figures comparing performances of all GRSs of all investigated traits in the WHI Hispanics, as well in the WHI African American (secondary analysis).

## Acknowledgements

The authors thank the staff and participants of HCHS/SOL for their important contributions. The Hispanic Community Health Study/Study of Latinos is a collaborative study supported by contracts from the National Heart, Lung, and Blood Institute (NHLBI) to the University of North Carolina (HHSN268201300001I / N01-HC-65233), University of Miami (HHSN268201300004I / N01-HC-65234), Albert Einstein College of Medicine (HHSN268201300002I / N01-HC-65235), University of Illinois at Chicago - HHSN268201300003I / N01-HC-65236 Northwestern Univ), and San Diego State University (HHSN268201300005I / N01-HC-65237). The following Institutes/Centers/Offices have contributed to the HCHS/SOL through a transfer of funds to the NHLBI: National Institute on Minority Health and Health Disparities, National Institute on Deafness and Other Communication Disorders, National Institute of Dental and Craniofacial Research, National Institute of Diabetes and Digestive and Kidney Diseases, National Institute of Neurological Disorders and Stroke, NIH Institution-Office of Dietary Supplements. The Genetic Analysis Center at the University of Washington was supported by NHLBI and NIDCR contracts (HHSN268201300005C AM03 and MOD03). Funding support for the “Epidemiology of putative genetic variants: The Women’s Health Initiative” study is provided through the NHGRI grants HG006292 and HL129132. The WHI program is funded by the National Heart, Lung, and Blood Institute, National Institutes of Health, U.S. Department of Health and Human Services through contracts HHSN268201100046C, HHSN268201100001C, HHSN268201100002C, HHSN268201100003C, HHSN268201100004C, and HHSC271201100004C. The authors thank the WHI investigators and staff for their dedication, and the study participants for making the program possible. A full listing of WHI investigators can be found at: https://www.whi.org/about/SitePages/Study%20Organization.aspx. T.S. was supported by the NHLBI [R01 HL120393-03S1 and 1R35HL135818]. K.G. was supported by the National Science Foundation Graduate Research Fellowship Program under Grant No. DGE-1256082. Any opinions, findings, and conclusions or recommendations expressed in this material are those of the author(s) and do not necessarily reflect the views of the National Science Foundation.

